# QC4Metabolomics: Real-time and Retrospective Quality Control of Metabolomics Data

**DOI:** 10.1101/2024.12.29.630653

**Authors:** Jan Stanstrup, Lars Ove Dragsted

## Abstract

**Motivation:** The ability to answer complex biological questions in metabolomics relies on the acquisition of high-quality data. However, due to the complex nature of liquid chromatography–mass spectrometry acquisition, data quality checks are often not done comprehensively and only at the post-processing step. This can be too late to mitigate analytical problems such as loss of *m/z* calibration, retention time drift and severe ion suppression. It is often not practically or economically feasible to reanalyze samples, and interpretation of the acquired compromised data, if at all possible, is limited, despite the considerable expenses incurred to obtain them.

**Results:** We therefore introduce `QC4Metabolomics`, a real-time quality control monitoring software for untargeted metabolomics data. `QC4Metabolomics` monitors files as they are acquired or retrospectively by tracking any user-defined compound(s) and extracting diagnostic information such as observed *m/z*, retention time, intensity and peak shape, and presents the results on a web dashboard. `QC4Metabolomics` also monitors the levels of common or user-defined contaminants. We report herein real-world examples where `QC4Metabolomics` easily reveals analytical problems retrospectively that could have been immediately addressed with real-time monitoring, so that the samples would have been analyzed without any quality control issues.

**Availability and Implementation:** `QC4Metabolomics` is available as code at https://github.com/stanstrup/QC4Metabolomics. A `Docker` image, a `docker-compose` setup file and demo data are also provided for easy deployment.

## Introduction

Metabolomics studies aim at the simultaneous determination of many compounds in complex matrices. Their success relies greatly on appropriate experimental design, acquisition of high-quality data and suitable data (pre)-processing. The latter two determine the ability to identify and reveal the relative amounts of each single metabolite in these complex mixtures which help answer the biological question. In contrast to other types of analyses, for untargeted mass spectrometry (MS)-based metabolomics, it is very difficult to fully assess the quality of the data due to its complexity.

Several kinds of samples are typically added in an untargeted sample analysis sequence to assess the performance of the analytical instrument. The added samples typically include a pool of all study samples, a so-called quality control (QC) sample. Assay blanks and other “blank” injections and potentially mixed standards or other samples used for quality assessment and diagnostic purposes are typically also used^1,2^. The QC sample is injected at regular intervals, i.e. every 5-10 injections among the biological samples. Isotopically labelled compounds, so-called internal standards, are commonly added to all samples, even if they do not represent specific analytical targets. The operator will ordinarily attempt to use the pooled QC samples to assure the data quality during acquisition by spot-checking the internal standards or other common metabolites for known problems that can arise when analyzing a long sequence of samples.

The pitfall of relying on manual spot checking is that many issues exist that are not obvious, such as loss of calibration in a limited mass region, non-obvious contaminants, ion suppression in specific chromatographic regions or ion suppression or enhancement affecting only certain peaks. Many of these critical problems often remain unnoticed until the data analysis stage. At this stage, the data analyst is left attempting mathematical corrections to the data, which may help but this is not always sufficient. Broadhurst et al. and González-Domínguez et al. have reviewed such procedures as well as other good practices for using QC samples^3,4^. The software `QCScreen`^5^ gives a good overview of quality and stability parameters and `QC-MXP`^6^ provides tools for correcting intensity drifts. These programs allow post-analysis corrections, but not real-time diagnostics and a laboratory could therefore end up with a situation where hundreds of samples have been analyzed at considerable costs, only to realize at the *data analysis* stage that the data needs extensive post-hoc modification or even that the quality is irreparably compromised.

Here, we present `QC4Metabolomics`, an open-source Shiny web application and R package that monitors QC parameters in metabolomics experiments in (near) real-time and documents data quality post-analysis using an intuitive dashboard. We believe that `QC4Metabolomics` would enable users to spot and correct for analytical problems during analysis, thus ensuring the acquisition of the highest quality data.

## Experimental Section

### Installation and Data Extraction

`QC4Metabolomics` leverages the rich ecosystem of R packages. A review of metabolomics-related R packages has been published previously^7^. The entire system is containerized with `Docker`^8^ for easy installation and portability and settings are changed by passing environmental variables to the `Docker` container.

Data pre-processing is primarily done using `xcms`^9^ and other packages under the “RforMassSpectrometry” initiative^10^. The graphical user interface (GUI) is built with `Shiny`^11^ and interactive plots are created using `ggplot2`^12^ and `plotly`^13^ with `viridis` color gradients^14,15^. Data is stored in a `mariaDB`^16^ and all required R packages are managed with `renv`^17^.

### Data Transfer and Format Conversion

In `QC4Metabolomics` two optional helper programs are provided for data export and conversion.

1. A Windows batch script that is intended for use on the instrument PC to copy files to e.g. a network drive that `QC4Metabolomics` can access. While this script is specific to Waters’ “.raw” files the rest of the program is agnostic to the original vendor format. The script will move files named according to a given pattern to a new location when the run is completed but also make symbolic links from the original location to the destination. This ensures that the files can still be accessed from the vendor software but are typically also available on a storage server that `QC4Metabolomics` can then access. The destination file path of the file is written to a text file so that it can be found by down-stream processes.
2. An automatic converter that uses `ProteoWizard`^18,19^ (in `Docker` and thus OS-agnostic) to convert vendor-specific raw data to mzML. It reads the text file with the file paths of vendor files and converts them such that the mzML files are available for further analysis by `QC4Metabolomics`. The destination path of the mzML is written to a text file that `QC4Metabolomics` can read to detect new files. These systems can be modified as needed for the specific setups.

### QC4Metabolomics Modules

`QC4Metabolomics` itself consists of two independent processes. A continuously running process for processing and analyzing new data, paired with a Shiny web server that delivers an interactive graphical dashboard, enabling real-time user interaction and monitoring. A database collects the data generated in the first process to make it available for the second process. The workflow is illustrated in Figure 1.

**Figure 1:**
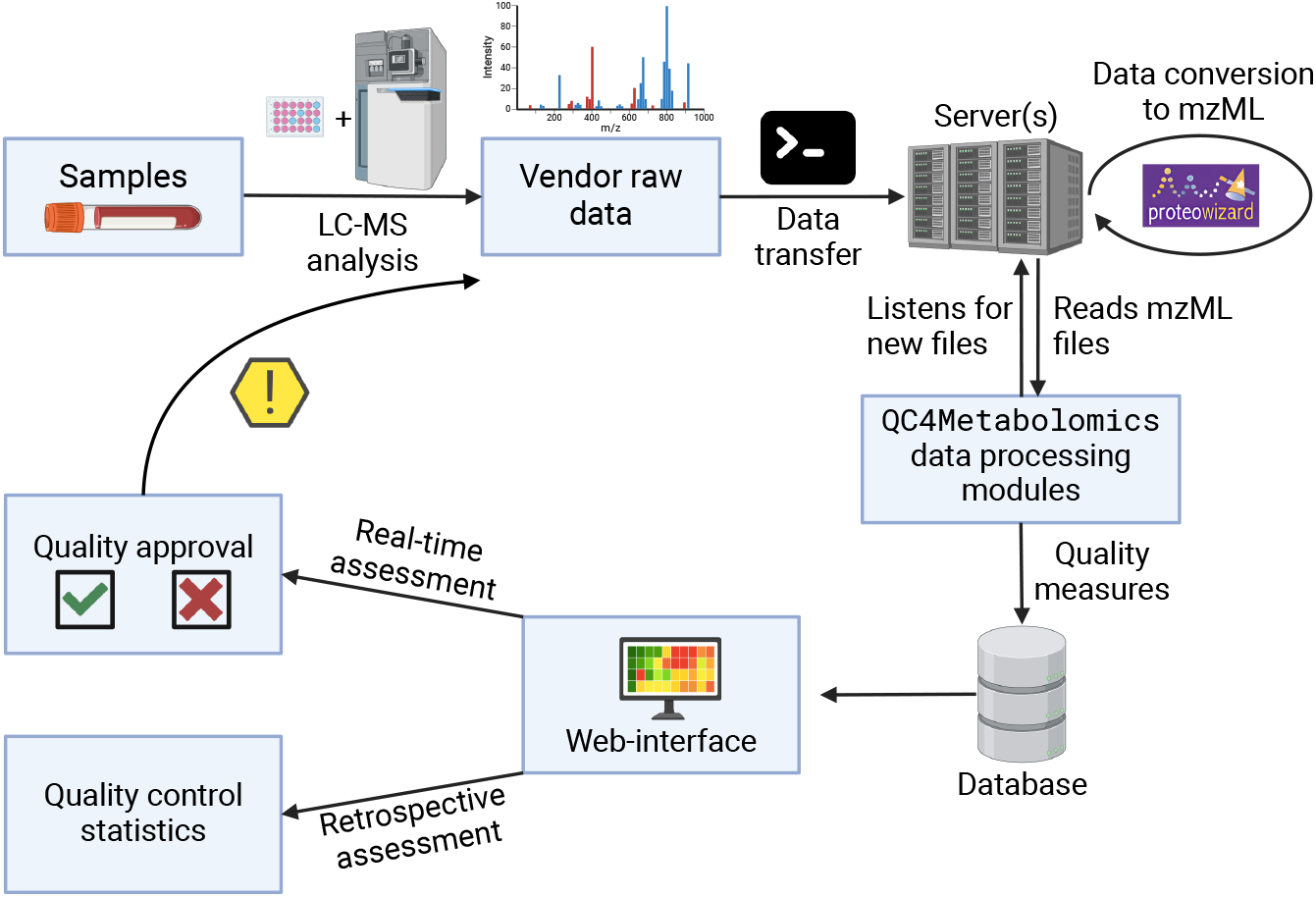
Workflow of `QC4Metabolomics`. When samples are analyzed by the LC-MS system the raw data is transfered by a simple batch script to a server (optional) accessable by `QC4Metabolomics`. The data is then automatically converted to mzML by MSconvert from `ProteoWizard`. If new mzML files exist, this triggers `QC4Metabolomics` to start processing the files and write the resulting quality parameter data to a database. This data is then available on a web-interface where the user in real- time can decide whether or not the system is sufficiently performant and take actions as neccesary. Retrospectively the data can also be reviewed for possible issues to be addressed during pre-procesing. Created in BioRender. Stanstrup, J. (2024) https://BioRender.com/w48h302.

`QC4Metabolomics` is also structured in modules performing specific data analyses during the data analysis process or adding new GUI elements to the dashboard. An R/Shiny programmer can add new modules without modifying existing `QC4Metabolomics` code such that each laboratory can adjust the system to their needs. There are currently 9 available modules.

1. `module_Files` looks for newly acquired files (mzML, mzXML, mzData etc.) either by traversing a directory or by reading from a text containing file paths and then adds them to a table in the database.
2. `module_File_schedule` checks if new filenames were added to the database and schedules them for processing by each enabled module that extracts data from the files (e.g. `module_File_info`, `module_TrackCmp` and `module_Contaminants`).
3. `module_File_info` extracts information (e.g. instrument name, project name, ionization mode and sample name) from the filenames using a user-defined naming pattern and the precise analysis time from the mzML files’ metadata.
4. `module_TrackCmp` detects peaks based on a user-defined list of *m/z* and retention time (RT) pairs (separately for different instruments, see Figure S1). The GUI displays diagnostics (e.g. *m/z* and RT deviation, peak shape and intensity) on a time-line for easy monitoring of systematic changes. The displayed analyses can be filtered by keywords and regular expressions.
5. `module_Contaminants` uses a list of nearly 800 known contaminant ions^20,21^ or a user-supplied list. The module offers three ways to monitor contaminants, as outlined in the Results section below.
6. `module_Productivity` provides a calendar overview of which projects were run and the number of injections run each day. See Figure S2.
7. `module_Log` displays a log of the activities of the other modules. See Figure S3.
8. `module_Debug` provides technical information on the `QC4Metabolomics` setup. See Figure S4.

Modules can be enabled or disabled and settings can be adjusted by passing environmental variables to the `Docker` container.

## Results and Discussion

`QC4Metabolomics` transforms the tedious manual spot-checking into an automatic process where the operator chooses a selection of analytes to be monitored for various quality parameters in real-time. These analytes could be internal standards or any common analyte and ideally of different structural classes and with RTs spanning the duration of the chromatographic runs and mass range. The operator will monitor the performance of the instrument as analyses are being done or use the data collected to compare present and previous analytical series. This fills a gap in the untargeted metabolomics methodology to improve QC and avoid wasting machine time, resources and irreplaceable biological material on suboptimal analyses.

### Monitoring of Retention Time Shifts

Figure S5 shows the web app’s structure. The upper panel allows access to different modules and selection of individual instruments. The body displays the content of the selected module (here `module_TrackCmp`). The database can be queried by one or more projects, ionization mode, sample ID, and filtered by analysis date. The screenshot also shows the RT and *m/z* deviation data for our long-term QC pool (MetNexs) analyzed within all batches, and we can observe that in the shorter term the deviations are normally less than 0.05 min, while the differences between different batches can be at least twice as large.

In addition, peak shape (tailing factor and asymmetry factor) can also be monitored to indicate column degradation (see Figure S6).

### Monitoring of Mass Accuracy and Calibration

Different vendors handle calibration differently during acquisition. Some inject a calibrant between injections, while others, like the Waters Synapt instrument used for the examples in this paper, calibrate during analytical runs. The instruments will typically not alert the user or stop the acquisition if the calibration is inaccurate. The calibrant used may cover the analyzed mass range incompletely and higher *m/z* deviation might be observed for some mass ranges. Monitoring several compounds across the relevant *m/z* range in `QC4Metabolomics` would allow the operator to realize such problems in real time.

In Figure 2 we observe instances of suboptimal calibration. In May/June of 2022 a small systematic offset was observed. This is unlikely to be a practical issue but in June, sensitivity decreased by an order of magnitude which in turn reduced precision. At the end of 2022 and beginning of 2023, the calibrant signal was lost intermittently, then completely, compromising calibration. It was later confirmed that the pump responsible for the flow of calibrant was malfunctioning and thus the calibration signal was too low to be used reliably for calibration. These are examples where routine use of `QC4Metabolomics` could have identified critical issues early allowing for timely intervention before more samples were analyzed.

**Figure 2:**
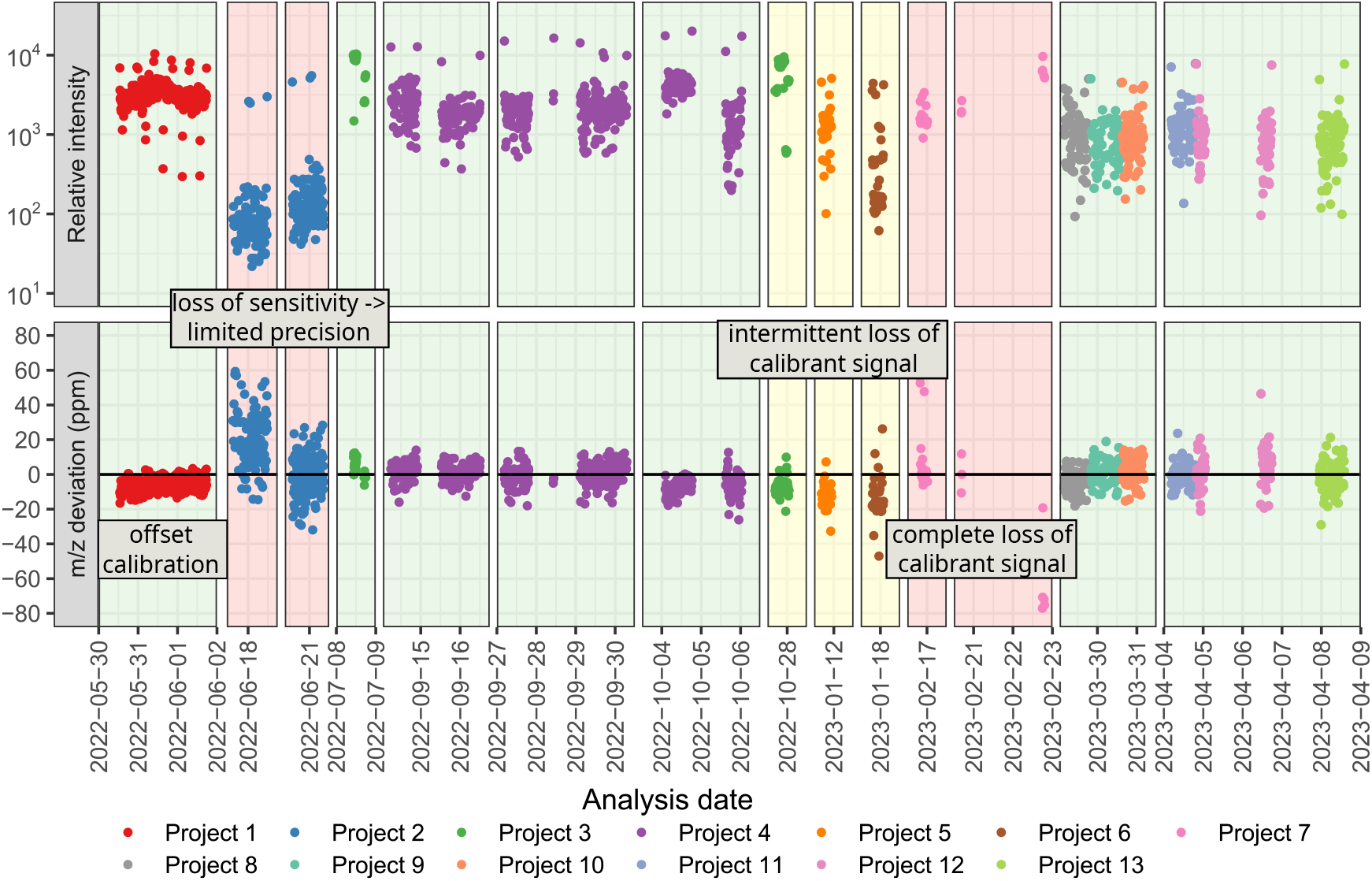
Example of monitoring performance with `QC4Metabolomics`. All injections between May 2022 and April 2023 on a single instrument are shown using the ion corresponding to tryptohan as the example. The upper panel shows the intensity on a log scale while the lower panel shows the relative *m/z* deviation. The time scale has been trimmed to exclude time periods where the instrument was not in use for metabolomics analyses. The plot is not a screenshot from `QC4Metabolomics` but constructed based on data extracted from our `QC4Metabolomics` database.

### Monitoring of Intensity Drift

Loss of intensity in the mass detector is often due to matrix build-up on the ion source sprayer. This can be monitored post-acquisition by looking at systematic drifts in a principal component analysis (PCA) scores plot^22^. Here, we are instead monitoring specific ions, to assess their drift in real-time. A drop in sensitivity during the analysis of a batch of “dirty” biological samples (e.g. serum, fecal extracts) is unavoidable, but a large drop indicates introduction of new contaminants, or that matrix build-up has reached unacceptable levels^23^.

In Figure S7 we can see how the intensity has decreased during a three-week period for tryptophan measured in our long-term QC samples. We can clearly see both intra- and inter-batch effects using this plot. It is then up to the operator to assess if the variation is acceptable or if action needs to be taken.

### Monitoring Contaminants

Contaminants can severely compromise the validity of an analysis, and they can be very difficult to detect manually, since they may not even form peaks but add variable levels of background noise. The introduction of contaminants to the mass analyzer can stem from several sources including the solvents, column bleed and matrix build-up causing ion suppression and irrelevant features^24^. Depending on the source of the contaminant, they may show up as additional peaks (i.e. they have been retained by the column) or as a persistently present mass covering the whole chromatogram (post-column contamination or not retained). These “additional” peaks can be monitored manually if they are sufficiently large to show up in the total ion chromatogram (TIC). However, if the peaks are relatively small this becomes impractical and persistently present masses can only be detected if they have an intensity large enough to significantly change the baseline intensity. `QC4Metabolomics` uses a list of nearly 800 known contaminant ions^20,21^ or a user-supplied list and offers three ways to monitor contaminants:

#### 1. Time view

Plots the changes over time for a single contaminant. This can give insights into the source of the contaminant e.g. by realizing that blank samples do not contain the contaminant. See Figure S8.

#### 2. Sample composition

Intensity-sorted barplot of contaminants in a single sample to easily determine major contaminants. See Figure S9.

#### 3. Heatmap

A heatmap showing analysis time on the x-axis and clustered contaminants on the y-axis, visualizing patterns and changes over time to help identify the sources. See Figure S10.

For all 3 plots the user can select between using the max intensity of the extracted-ion chromatograms (EICs), representing good estimates of peak height for peak-like contaminants (e.g. nylon), or the mean intensity across the EICs, which gives a more accurate picture of contaminants that have a consistent presence (e.g. polyethylene glycols (PEGs)) across the whole RT range. Once presence of a severe contaminant have has been realized, it is up to the operator to assess the impact on the analyses and take actions to track down the source of the contaminantion.

A similar approach to the one presented here was first described for the processing tool `XCMS online`^25^, however it is currently not available and not open-source. `Rapid QC-MS`^26^, a web dashboard that provides real-time quality metrics similar to `QC4Metabolomics` was recently reported. However, `Rapid QC-MS` is focused on tandem MS experiments and monitoring specific reference samples as opposed to `QC4Metabolomics` that monitors all samples and is focused on untargeted full-scan MS experiments.

We envision extending `QC4Metabolomics` to create static project reports summarizing quality metrics and additional analyses. E-mail notification to operators of aberrant readings could also be implemented, but defining threshold values remains difficult for most metrics. However, consistent use of `QC4Metabolomics` can supply the data needed to establish reasonable thresholds. For example the data in Figure 2 indicates that deviations larger than 20 ppm would indicate serious analytical problems that should be mitigated.

## Conclusion

`QC4Metabolomics` provides a robust, real-time and retrospective QC monitoring system. This proactive approach helps identify and rectify analytical issues at an early stage and thus facilitates better data quality, conserving valuable resources and enhancing the reliability of the study outcomes.

The easy-to-use GUI provides an intuitive and interactive experience for users.

Furthermore, the use of `Docker` for deployment ensures ease of installation and portability across different computational environments while the modular system makes it easy to customize to specific needs.

With broad community adoption, we envision `QC4Metabolomics` as a key safeguard against poor data quality, contributing to more robust datasets and efficient resource use.

## Data Availability Statement

The Shiny app is available as open-source code at https://github.com/stanstrup/QC4Metabolomics. A `Docker` image and a docker-compose setup file are also provided for easy deployment, along with a demo and sample data. The current version at the time of publication has been deposited at ZENODO with DOI: 10.5281/zenodo.14524880.

## Author Contributions

The features of the software were conceived by JS and LOD. The software was programmed by JS. The manuscript was first drafted by JS and contributed to by LOD. All authors have given approval to the final version of the manuscript.

## Acknowledgements

We thank Sarah Fleischer Ben Soltane for sample analysis, feedback and testing and Giorgia La Barbera for manuscript suggestions and proof-reading. This project was funded by grants from the Carlsberg Foundation (Semper Ardens, CF15-0574) and Novo Nordisk Foundation Challenge programme (PRIMA, grant number NNF19OC0056246).

