## Supplemental figures for "QC4Metabolomics: Real-time and Retrospective Quality Control of Metabolomics Data"

### Supplementary Material for QC4Metabolomics: Real-time and Retrospective Quality Control of Metabolomics Data

29. December 2024

[1] Department of Nutrition, Exercise and Sports, University of Copenhagen, Rolighedsvej 30, 1958 Frederiksberg

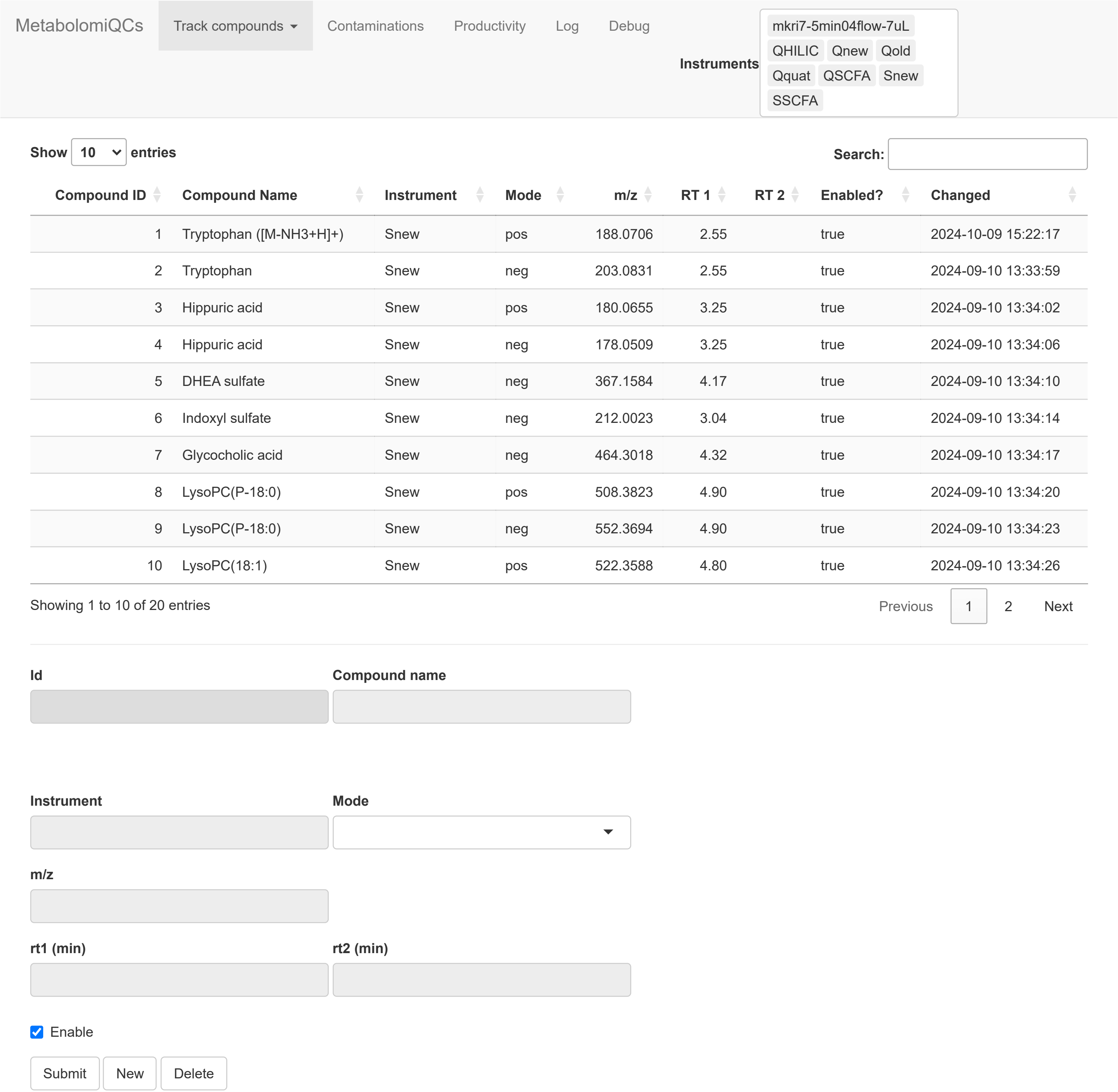


Figure S1: Screenshot of `module_TrackCmp` for the settings for the tracked compounds. New compounds can be added or retention time and m/z adjusted for future analyses.

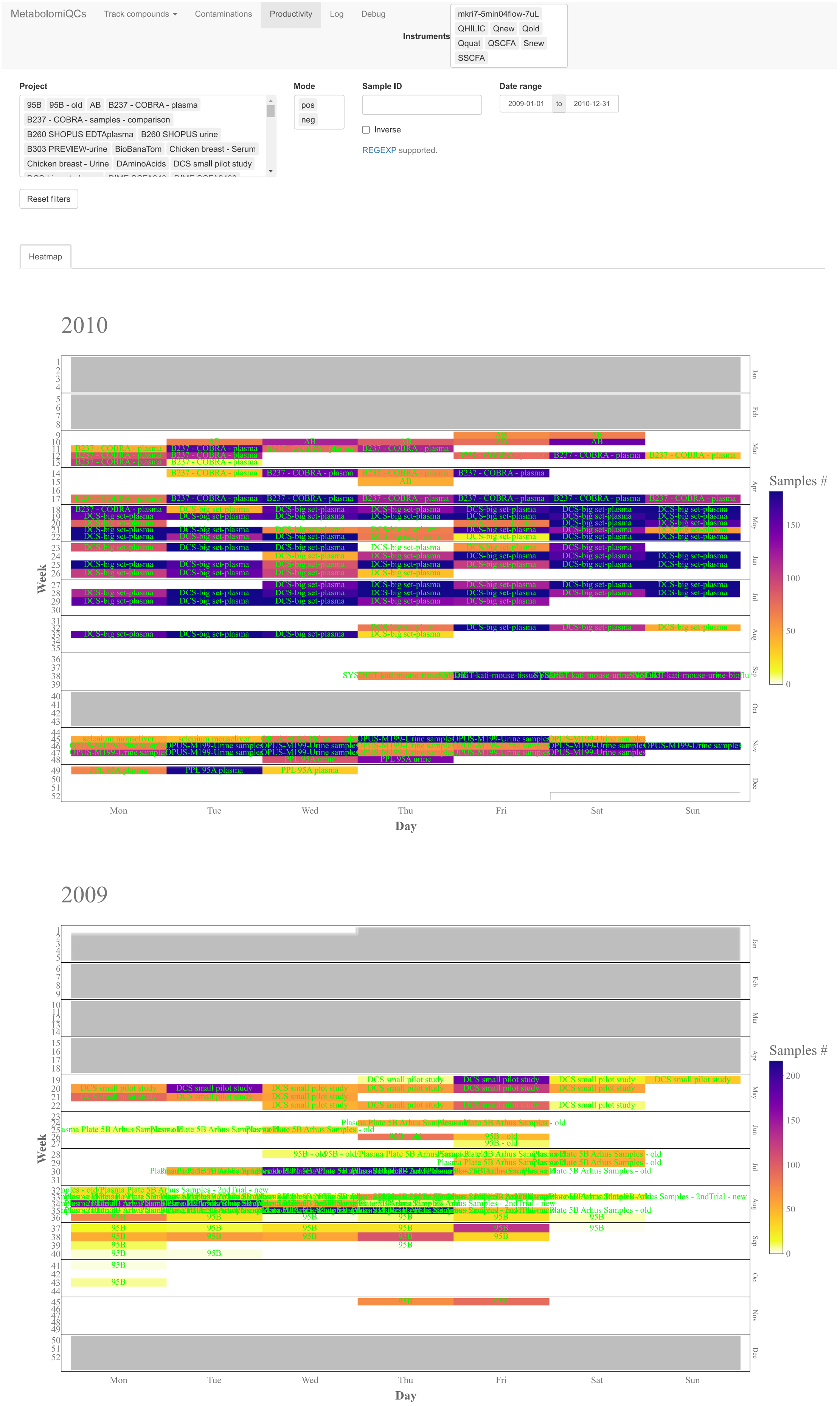


Figure S2: Screenshot from `module_Productivity` showing an overview of projects analyzed in each month. The color indicates the number of samples.

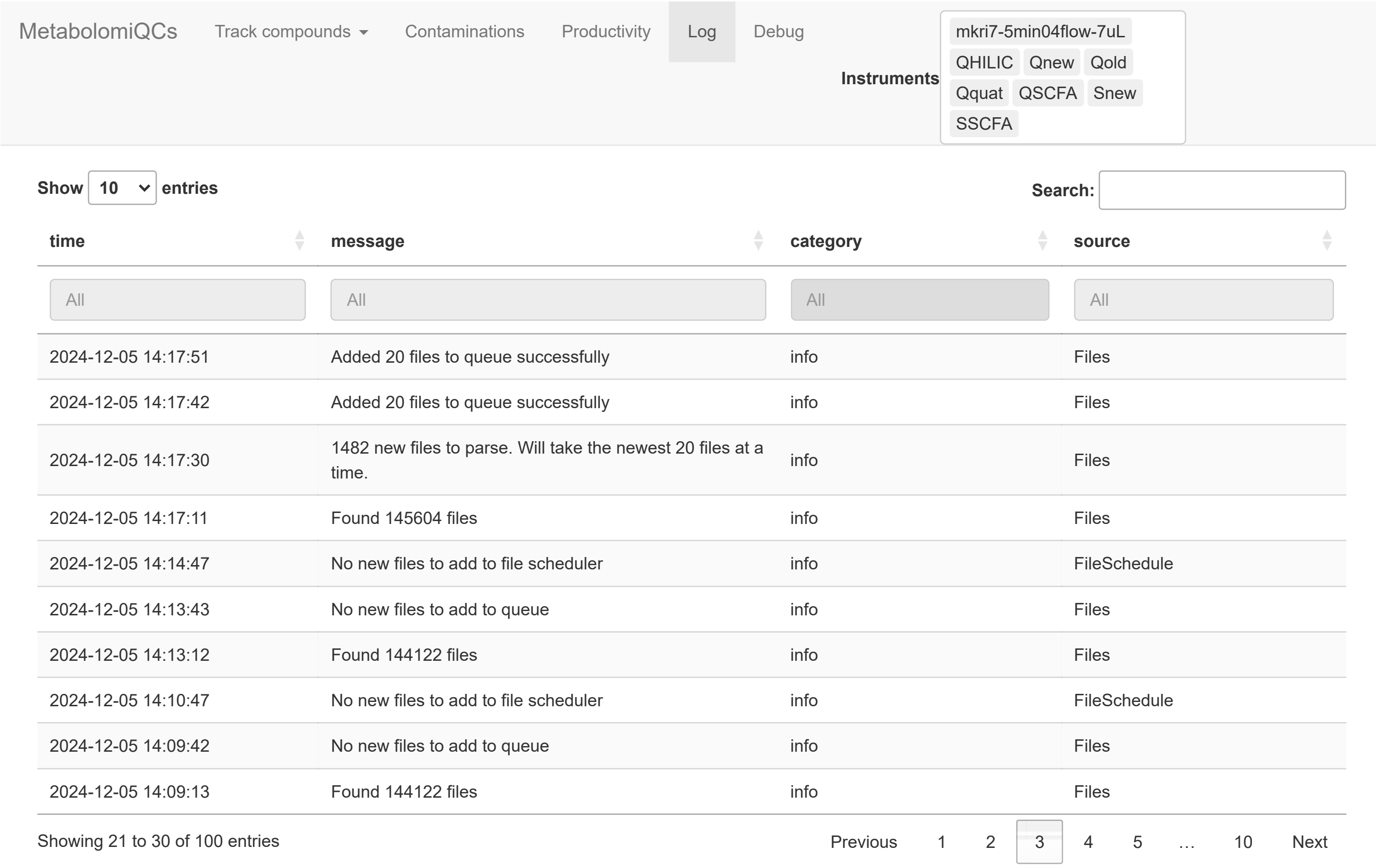


Figure S3: Screenshot of `module_Log`. Time-stamped activity can be read and filtered.

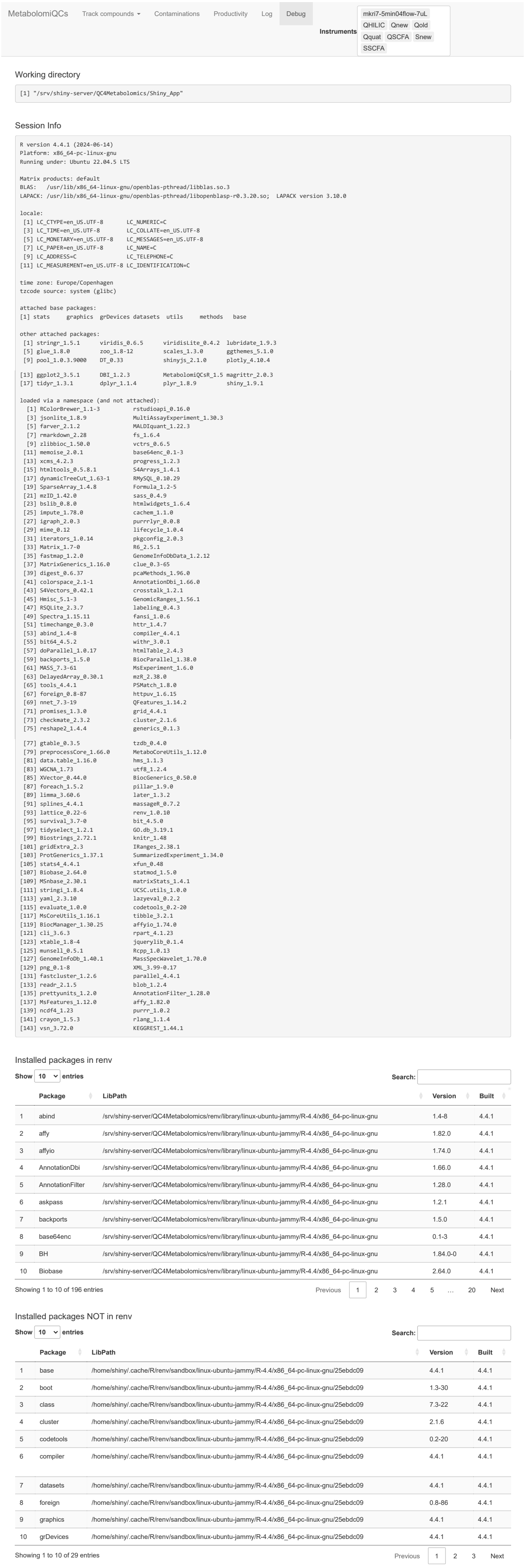


Figure S4: Screenshot of `module_Debug`. The R and R package versions are shown as well as the location of R packages. This module is meant only for development debugging.

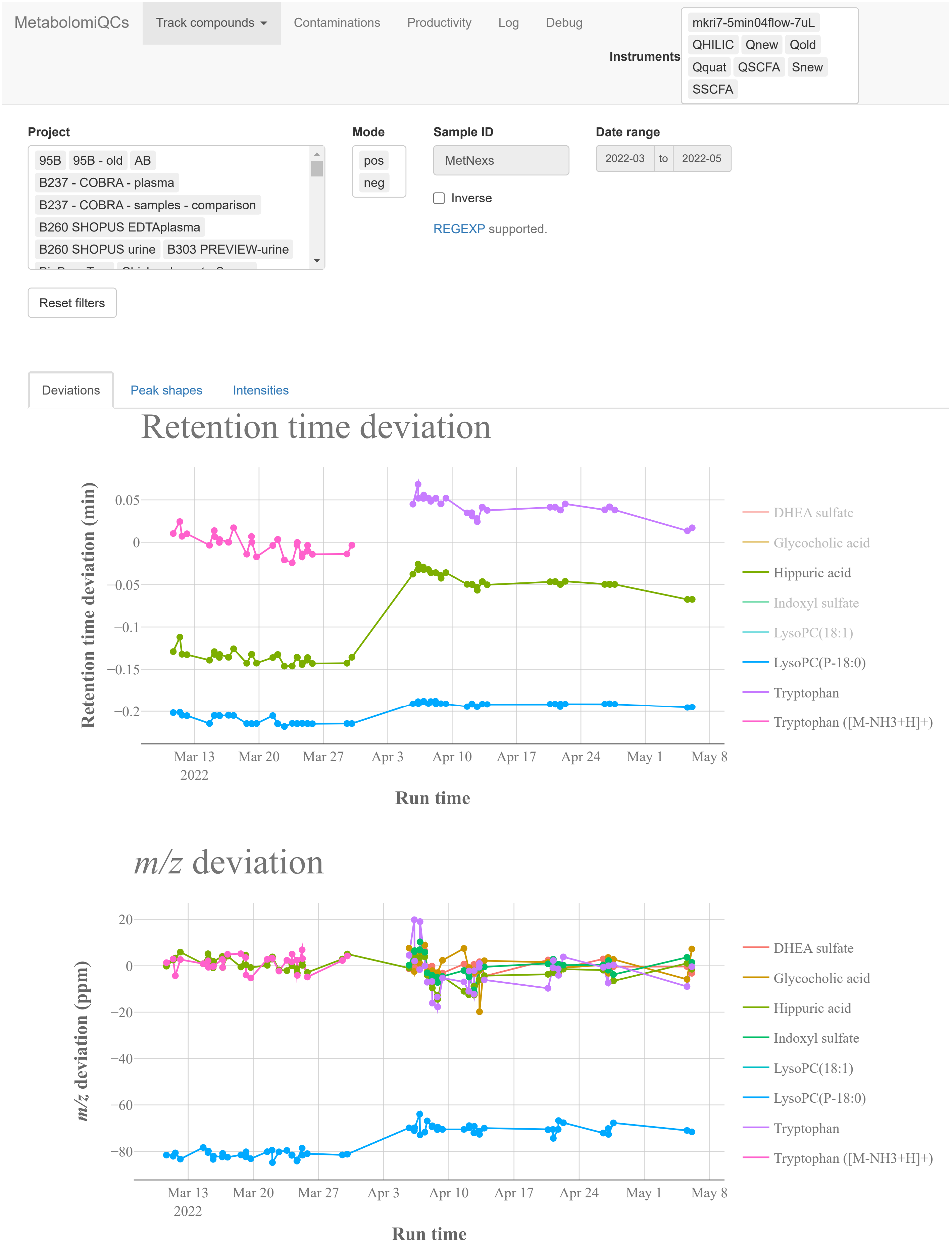


Figure S5: Screenshot of `module_TrackCmp` showing the retention time and $m/z$ deviation from the expected values for samples labelled "MetNexs" which is our own longterm reference material of human blood. In March positive mode was analyzed and in April negative mode was analyzed. Four observations can be made. 1) Only for Hippuric acid is the expected retention time in line with the observed retention time, the rest are offset somewhat. This is usually not a practical problem if a difference is only present between different projects. 2) The $m/z$ is far off for LysoPC(P-18:0) indicating that a different analyte have been picked by the system. 3) After the break in analyses in the beginning of April the retention times systematically increased, which should be investigated. 4) Apart from LysoPC(P-18:0) the $m/z$ was stable, but more so in positive mode.

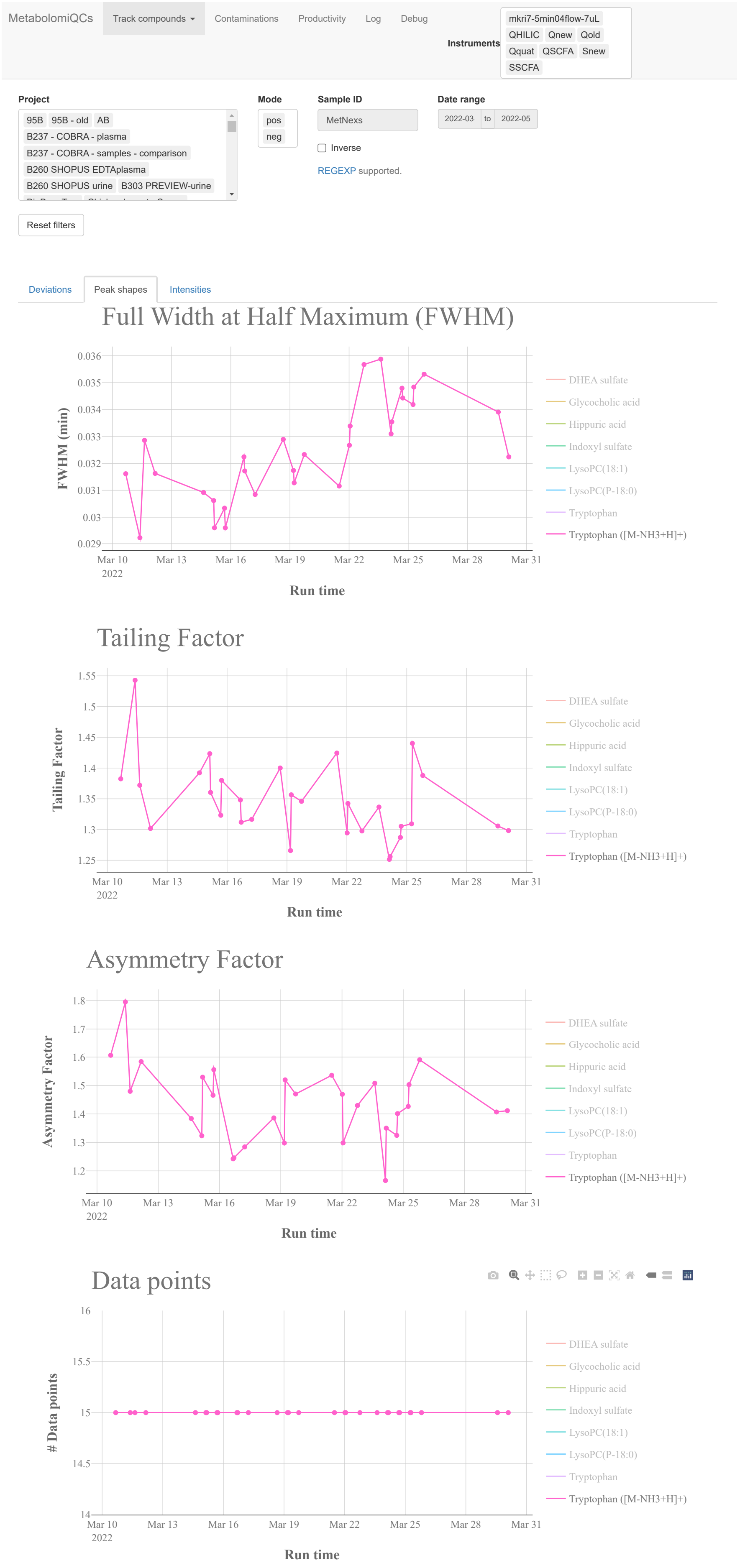


Figure S6: Screenshot of `module_TrackCmp` and as in Figure S7 Tryptophan is traced. The peaks' Full Width at Half Maximum (FWHM) indicates that peak broadening is occurring, while Tailing Factor, Assymmetry Factor and number of data points in the peak remains relatively constant.

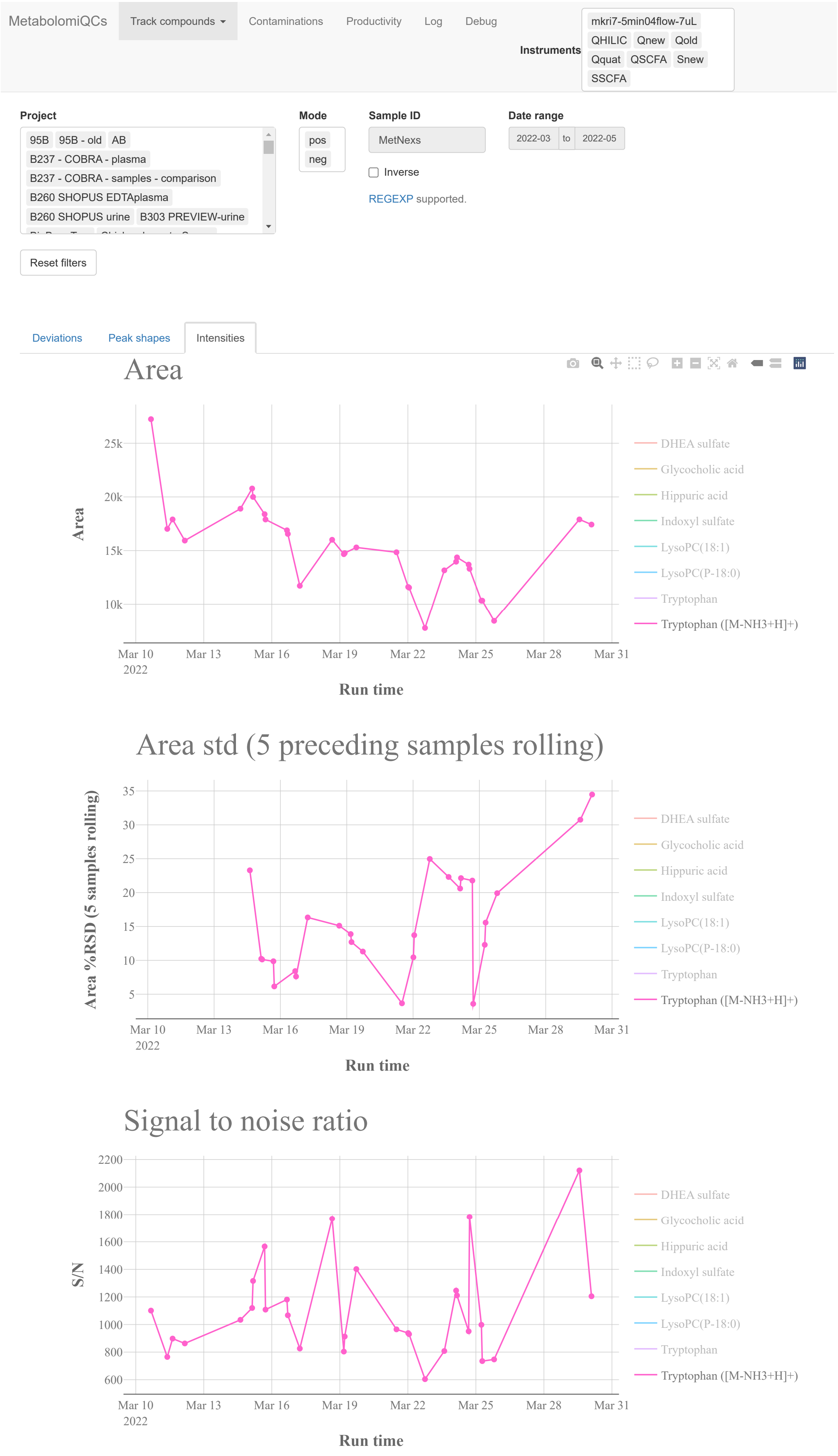


Figure S7: Screenshot of `module_TrackCmp` showing the same samples as in Figure S5 using Tryptophan in positive mode as the example. It can be observed that the intensity decreases over time to less than half the initial intensity. The RSD is consequently high.

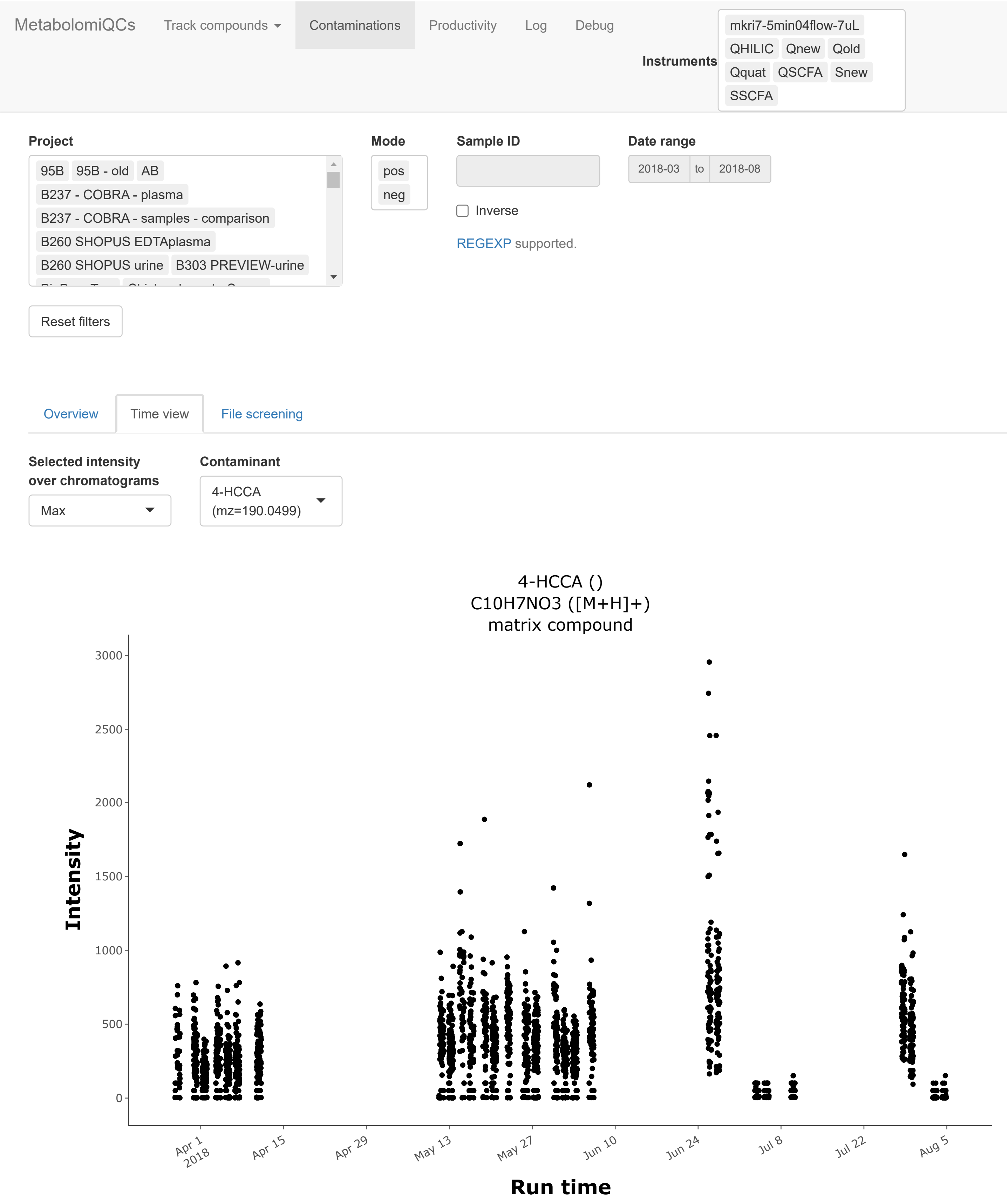


Figure S8: Screenshot of tracking of a specific contaminant with `module_Contaminants`. In the web GUI it is possible to hover over each point and see which sample a data point corresponds to. We can here realize that the samples with "zero" intensity for this contaminant are the blank samples and thus conclude that the contaminant comes from the sample preparation/handling rather than from the analytical system.

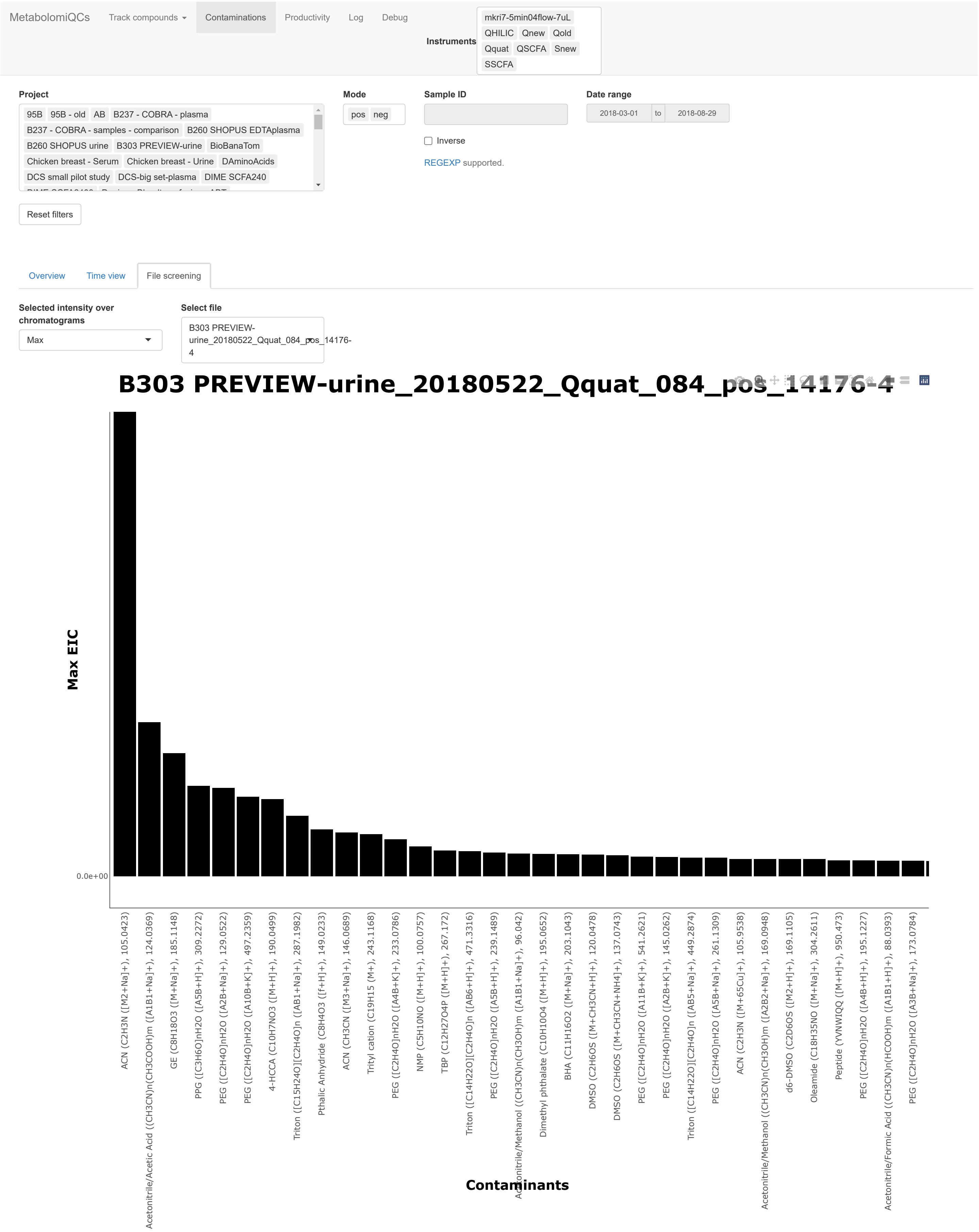


Figure S9: Screenshot from `module_Contaminants` of review of all detected contaminants in a single sample. The screenshot is zoomed to the most abundant contaminants.

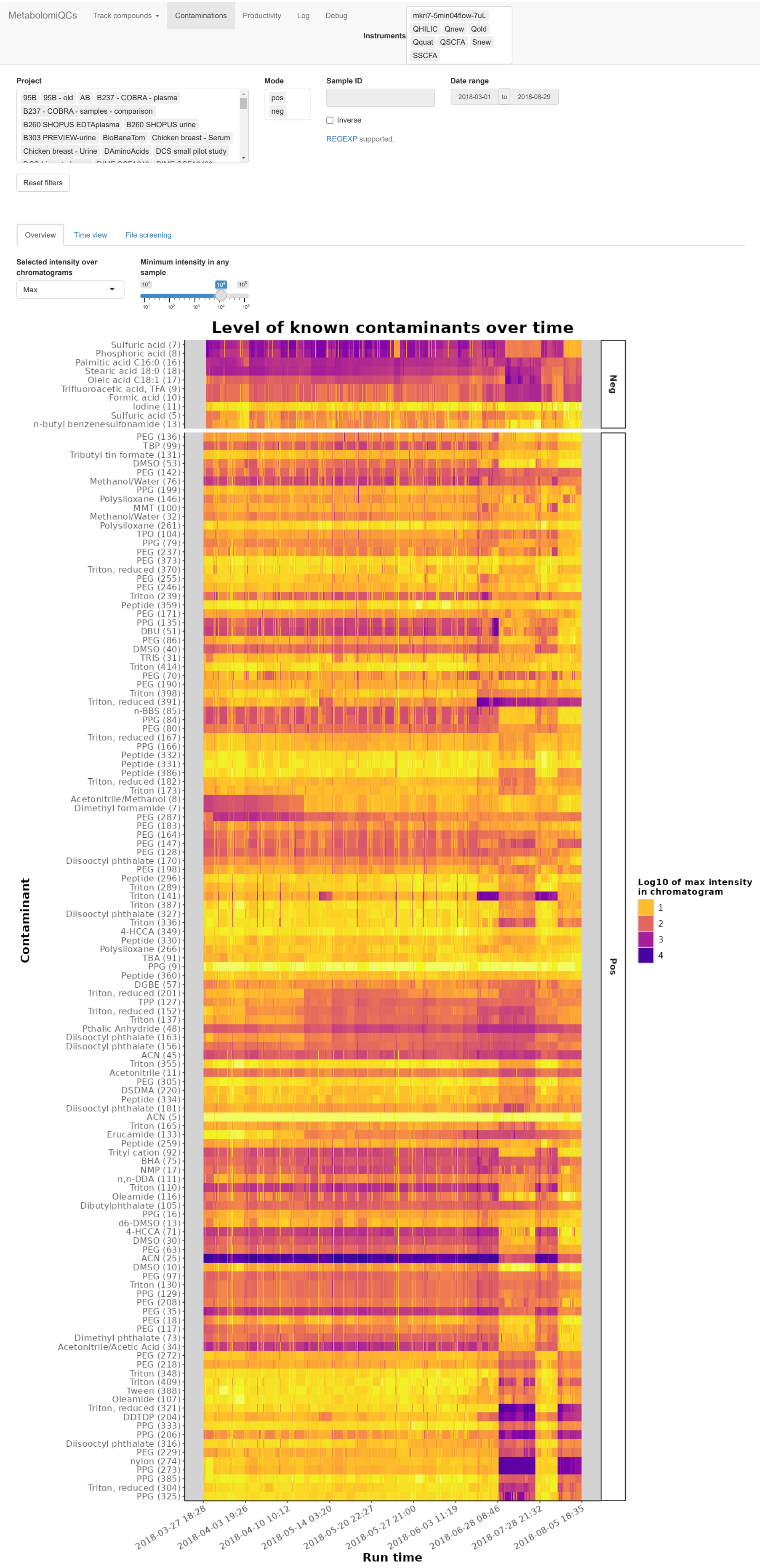


Figure S10: Screenshot of the overview heatmap from `module_Contaminants` showing the contaminants found in the samples. On the horizontal axis the detected contaminants are shown and on the vertical axis is the order the samples were analyzed in. The color scale indicates the intensity of the contaminant. The user can choose between plotting the maximum, mean or intensity found across scans in the samples. The heatmap tiles are drawn such that they start when the sample was analyzed and end when the next sample in the same ionization mode was started. Since there might be different amount of time between analyses that means that date and time is not equidistant in the plot. Data and time has been inserted along the axis to indicate approximately when a specific sample was analyzed.
